# Inhibition of the Dead Box RNA Helicase 3 prevents HIV-1 Tat and cocaine-induced neurotoxicity by targeting microglia activation

**DOI:** 10.1101/591438

**Authors:** Marina Aksenova, Justin Sybrandt, Biyun Cui, Vitali Sikirzhytski, Hao Ji, Diana Odhiambo, Matthew D. Lucius, Jill R. Turner, Eugenia Broude, Edsel Peña, Sofia Lizarraga, Jun Zhu, Ilya Safro, Michael D Wyatt, Michael Shtutman

**Affiliations:** Department of Drug Discovery and Biomedical Sciences, College of Pharmacy, University of South Carolina, Columbia, SC; Department of Statistics, College of Arts and Sciences, University of South Carolina, Columbia, SC; Department of Biological Sciences, College of Arts and Sciences, University of South Carolina, Columbia, SC; School of Computing, Clemson University, Clemson, SC

**Author notes:** To whom correspondence should be addressed. Corresponding authors: Correspondence to Michael Shtutman and Ilya Safro.

## Abstract

HIV-1 Associated Neurocognitive Disorder (HAND) is commonly seen in HIV-infected patients. Viral proteins including Tat cause neuronal toxicity and is worsened by drugs of abuse. To uncover potential targets for anti-HAND therapy, we employed a literature mining system, MOLIERE. Here, we validated Dead Box RNA Helicase 3 (DDX3) as a target to treat HAND via a selective DDX3 inhibitor, RK-33. The combined neurotoxicity of Tat protein and cocaine was blocked by RK-33 in rat and mouse cortical cultures. Transcriptome analysis showed that Tat-activated transcripts include makers and regulators of microglial activation, and RK-33 blocked Tat-induced activation of these mRNAs. Elevated production of proinflammatory cytokines was also inhibited by RK-33. These findings show that DDX3 contributes to microglial activation triggered by Tat and cocaine, and DDX3 inhibition shows promise as a therapy for HAND. Moreover, DDX3 may contribute to the pathology of other neurodegenerative diseases with pathological activation of microglia.

## Introduction

More than 36 million people worldwide are living with HIV infection and more than 1.2 million are in the USA based on World Health Organization (1). Neurological complications associated with HIV infection have long been known (2). With the introduction of highly active antiretroviral therapy (HAART), the life span of HIV-infected individuals has increased significantly to an average of 65 years (3). The increased life expectancy of the HIV-positive population is increasing the burden of the neurological complications to these patients and society. HIV-Associated Neurocognitive Disorder (HAND), a very concerning HIV-associated dementia, is prevalent in up to half of HIV-infected individuals and constitutes a growing health hazard in the aging population (4, 5). The risk of HAND and its associated neuropathology is higher among intravenous drug abusers (5, 6). The major drugs contributing to HIV pathogenesis are opiates and stimulants (cocaine and methamphetamine). Cocaine is the second most commonly abused drug in the US, and cocaine abuse in HIV-infected patients is associated with the worsening of HAND (7, 8).

During the early stage of infection, HIV enters the brain via trafficking of infected CD4^+^ cells and monocytes into the CNS (9). In the brain, HIV can infect macrophages, microglia, and astrocytes, with each of these as possible sites of persistence of latent HIV (10). Though neurons are refractory to viral infection, progressive neuronal damage has been observed in HIV-infected patients (10, 11). Direct HIV-mediated neurotoxicity can be caused by viral proteins that interact with neurons resulting in neuronal damage or death by multiple mechanisms, including disruption of calcium homeostasis, perturbation in glutamate and glycolytic pathways, inhibition of calcium and potassium channels, and others (10, 12, 13). Among these viral proteins, HIV-encoded Trans Activator of Transcription (Tat) continues to be synthesized and secreted by HIV-infected cells even in patients in which HAART therapy successfully prevents viral production (14). Tat uptake by uninfected cells results in both cytoplasmic and nuclear events promoting neuronal damage and death (15). The neurotoxicity of HIV and Tat is exacerbated by drugs of abuse, including opiates and cocaine. The combination of cocaine and Tat causes neurotoxicity that greatly exceeds that caused by cocaine or Tat alone (11, 16–19). Current HIV therapy options are focused on preventing entry of the virus into the brains of infected patients, but this therapy does not directly protect neurons and is not effective after entry has occurred. Thus, there is no approved therapy for the treatment of HAND and particularly for the combined neurological effects of HIV and drugs of abuse. Consequently, there is an urgent need to discover neuroprotective therapy that can alleviate HAND symptoms.

Artificial Intelligence (AI) – based discovery systems for biomedical literature analysis and hypothesis generation help researchers to navigate through the vast quantities of literature and can rapidly reveal hidden connections in the literature network (20, 21). Machine learning approaches can accelerate drug repositioning to discover new applications for existing compounds (22). We recently developed a new AI-based literature mining system, MOLIERE which is different from many other literature-based analysis systems (23, 24). The key difference is in the algorithmic pipeline and the amount of processed data. MOLIERE does not filter the data extracted from the papers, whereas most other systems work with only highly filtered semantic objects (such as MESH terms or only genes). In addition to our own ranking methods, the algorithmic pipeline relies on several recently developed scalable machine learning methods that have not been adopted by other knowledge discovery systems such as low-dimensional representation manifold learning and scalable probabilistic topic modeling (23, 24). In the present article, we performed MOLIERE analysis for possible links of human proteins with HAND in the biomedical literature. This analysis revealed a previously unknown connection between HAND and Dead Box RNA Helicase 3 (DDX3) (25, 26). DDX3 has not previously been associated with neurodegeneration related to HIV proteins, but was shown to be essential for translation of HIV proteins and the nuclear export of HIV RNA (27). Our results show that a known inhibitor of DDX3 called RK-33, which was originally developed and tested for anti-cancer therapy (28, 29), protects primary cortical neurons from neurotoxicity and inhibits the combined Tat plus cocaine dependent activation of microglia. The results validate DDX3 inhibition as a potential target for HAND therapy.

## Results

### Literature mining to uncover targets and small molecules to test for HAND therapy

To determine genes with unknown implicit connections to HAND, we utilized MOLIERE, a system to automatically generate biomedical hypotheses (23). The system builds a multi-modal and multi-relational network of biomedical objects extracted from Medline and ULMS datasets from the National Center for Biotechnology Information.(NCBI). In the current analysis we queried the list of all human genes downloaded from HUGO (30) paired with HIV associated neurocognitive disorder. The generated hypotheses were ranked based on a number of techniques described in (24). The hypotheses ranking represent the level of association each gene has with HAND. The genes were categorized as “Trivial” (top 2.5% genes with exceptionally high ranking scores typical of well-explored prior connections clearly associated with HAND), “High Potential” (next 15% of genes), “Low Potential” (next 15% of genes) and “Random” (Fig 1A). To determine the availability of small molecule inhibitors for “High Potential” category gene products, the genes were queried through a protein-small molecule interaction database, BindingDB (31). About 180 proteins out of 4450 genes have at least one associated small molecule (Fig 1B, Supplemental Table 1). Next, the list of the genes was narrowed by the selection of small molecule inhibitors that have been tested in animals and did not manifest any significant systemic toxicity at therapeutic concentrations. The search of the literature revealed 52 protein-small molecule pairs (Fig 1D, Supplemental Table 1). From the short list we selected RK-33, an inhibitor of Dead Box RNA Helicase 3 (DDX3) for experimental evaluation for the following reasons: RK-33 was shown to be a very selective inhibitor of DDX3 ATPase activity; the compound was active in vitro in nanomolar concentrations, and in micromolar concentrations in cell culture and animal models; and it was easily available commercially. Lastly, it had been extensively tested in rodents and did not show any systemic toxicity (28, 29). Although DDX3 had never been associated with HAND, it was shown to be important for HIV infection (27), specifically by exporting viral RNA from nucleus to cytoplasm (32).

**Figure 1.**
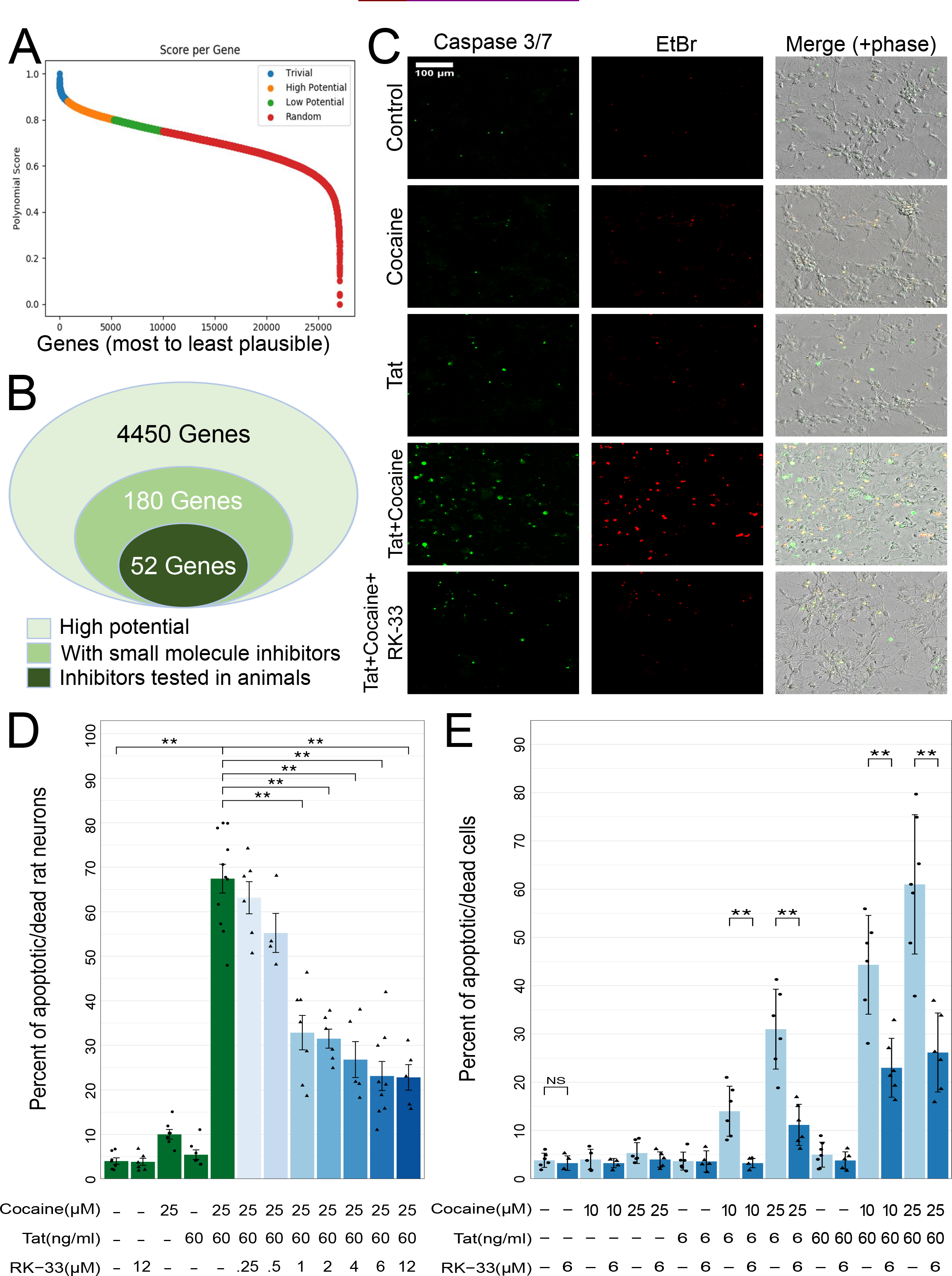
Selection and experimental validation of RK-33. The compound protects neurons in primary cortical cultures from the combined neurotoxicity of Tat and cocaine. Representation of ranking distribution of the ~27,000 HUGO genes. The ranked hypotheses representing the level of association of each gene with HAND were categorized as following: “Trivial” (top 2.5% genes with exceptionally high ranking scores typical of well-explored prior connections clearly associated with HAND), “High Potential” (next 15% of genes), “Low Potential” (next 15% of genes) and “Random” (**A**). Representative selection of genes for experimental validation, from all the “high potential genes” to genes with known small molecule ligands that have already tested safe for animal toxicity (**B**). Primary cortical cultures were treated with Tat (6 or 60 ng/ml) and/or cocaine (10 or 25 μM) or Tat combined with a range of RK-33 for 48 h prior to the addition of cocaine for another 24 h. Cultures were then fixed with 4% PFA and the dead/apoptotic cells were detected using CellEvent Caspase 3/7 Assay Kit (**C**: green, Caspase 3/7; red, Ethidium Bromide. Scale bar, 100 μm). Tat and/or cocaine only treatments are colored in green, and the deepening blue hue represents the increasing concentration of RK-33 (from 0.25 to 12 μM) (**D**). The bar graph shows Tat and/or cocaine only treatments in light blue and RK-33 in dark blue (**E**). The bar heights indicate mean values and error bars indicate one sample standard error from the sample mean. Each point corresponds to an image, with each image containing a range of 60-100 cells. The Mann-Whitney-Wilcoxon test is conducted to calculate the statistical significance, followed by Benjamini-Hochberg adjustment of p-values (*, p<0.05; **, p<0.01; ***, p<0.0001).

### RK-33 protects neurons in primary cortical cultures from the combined neurotoxicity of HIV-Tat and cocaine

To test the hypothesis that inhibition of DDX3 has neuroprotective effects in a model of HAND, we used a well-established cell culture model of rodent primary cortical neurons co-treated with Tat and cocaine in previously established concentration ranges (33–37). In this system, the addition of Tat at a low (6 ng/ml) or high (60 ng/ml) concentration, or cocaine (10 μM or 25 μM) individually did not significantly induce the death of primary neurons during 72 h of treatment (Fig 1C, D). However, pre-treatment with Tat for 48 h followed by the addition of cocaine for another 24 h (Fig 2A, B) drastically increased neuronal death, as measured by activated Caspase 3/7 signal and nuclear localization of ethidium bromide homodimer (Fig 1C, D). Strikingly, treatment with the DDX3 inhibitor RK-33 at 6 μM resulted in a 50%-80% reduction of neuronal death caused by Tat and cocaine co-treatment (Fig 1C, D). The initial 6 μM concentration of RK-33 selected was similar to that used previously (29). RK-33 protected rat neurons from Tat/cocaine toxicity in a dose dependent manner starting as low as 1 μM (Fig 1E), which was comparable with RK-33 bioactivity observed in other cell-based assays (29, 38). Nearly identical qualitative results were observed with a mouse primary neuron cortical culture regarding Tat/cocaine damage and RK-33 mediated protection from damage (Supplemental Figure S1). A notable quantitative difference was that rat primary neurons appear to be much more sensitive to cocaine relative to the mouse neurons, because a 10-fold lower cocaine dose was used to promote neuronal apoptosis in combination with Tat. Hence, the results in both rat and mouse cortical cultures show that DDX3 inhibition is neuroprotective against the combined insult of Tat and cocaine.

**Figure 2.**
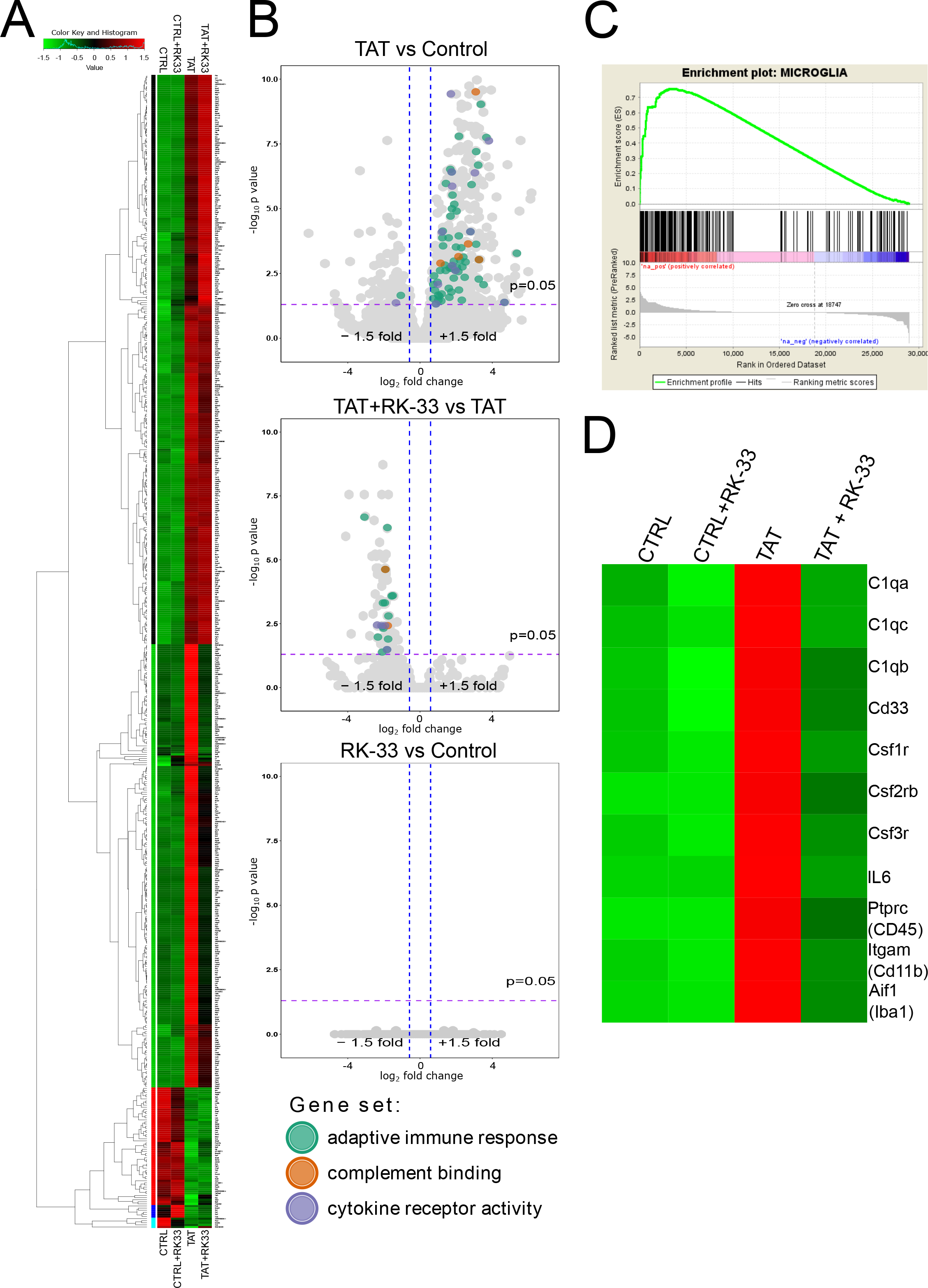
RNA-seq analysis transcriptome profiling of cortical cultures treated with Tat and RK-33. Cortical cultures were treated in triplicates with Tat (60 ng/ml), RK-33 (6 μM), combination of Tat (60 ng/ml) and RK-33 (6 μM) and then were compared with untreated cultures. RNA-seq transcriptomics profiling was performed as described (see Materials and Methods for details). Hierarchical clustering of log transformed counts per millions for genes differentially regulated by Tat treatment (p<0.0005) (**A**). Volcano plot comparing gene expression of control vs Tat-treated samples (upper panel), Tat plus RK-33 vs Tat-treated samples (middle panel), and control vs RK-33 treated samples (lower panel). Color indicates differentially expressing genes (FDR corrected p<0.05, absolute fold change >1.5) belonging to selected Enrichr GO categories/pathways (**B**). GSEA enrichment plot of Microglia-specific genes in Tat vs control dataset (**C**). Heatmap for expressions of selected genes representing the microglia cells (**D**).

### RNA-seq transcriptomic profiling of RK-33 effects on Tat-treated cortical cultures

To explore the mechanism of RK-33 neuroprotection seen above, we performed RNA sequencing (RNA-seq) of the primary cortical cultures treated with Tat in the presence or absence of RK-33. We found differential expression of 547 genes in Tat-treated cultures compared to control (Adjusted FDR, P<0.05). Hierarchical clustering revealed that Tat-dependent upregulation of 211 genes was inhibited by RK-33 (adjusted FDR P< 0.05 for 83 genes) (Fig 2A-B, Supplemental Figure S2, Supplemental Table 2). Strikingly, there was no statistically measurable effect on any gene expression changes caused by RK-33 treatment alone (Fig 2B, Supplemental Table S2). Pathway enrichment analysis of Tat-regulated genes shows that proinflammatory pathways, such as “Complement Cascade”, “Neutrophil Dysregulation” and “Cytokine Signaling” are significantly enriched, as noted by the comparisons between Tat vs control (Fig 2B, Supplemental Figure S3). In the major brain cell types (neurons, astrocytes, oligodendrocytes, epithelial and microglial cells), these pathways are known to be associated with activated microglia (39–41). Notably, the expression of these same genes was suppressed by RK-33 treatment (Fig 2B, Supplemental Figure S3). Among this subset of genes upregulated by Tat alone and sensitive to RK-33 the following are worth noting: complement components C1qa, C1qc, C1qb; bona fide markers of microglial cells such as ionized calcium binding adaptor molecule 1 (Iba1 or AIf1), integrin Cd11a/b (Itgam), Ptprc (CD45); Colony-stimulating factor-1 (CSF-1) and granulocyte colony‐stimulating factor (g-CSF) receptors, Csf1r and Csf3r, which are necessary for microglial survival and proliferation (42) (Fig 3D). Further, gene set enrichment analysis (GSEA) (43) with custom databases of genes enriched in brain cell subtypes as defined by RNA-seq Transcriptome and splicing databases of the cerebral cortex (44) shows significant enrichment of microglia-associated genes in Tat-regulated and RK-33 sensitive, Tat – regulated genes (Fig 2D, Supplemental Figure S4).

**Figure 3.**
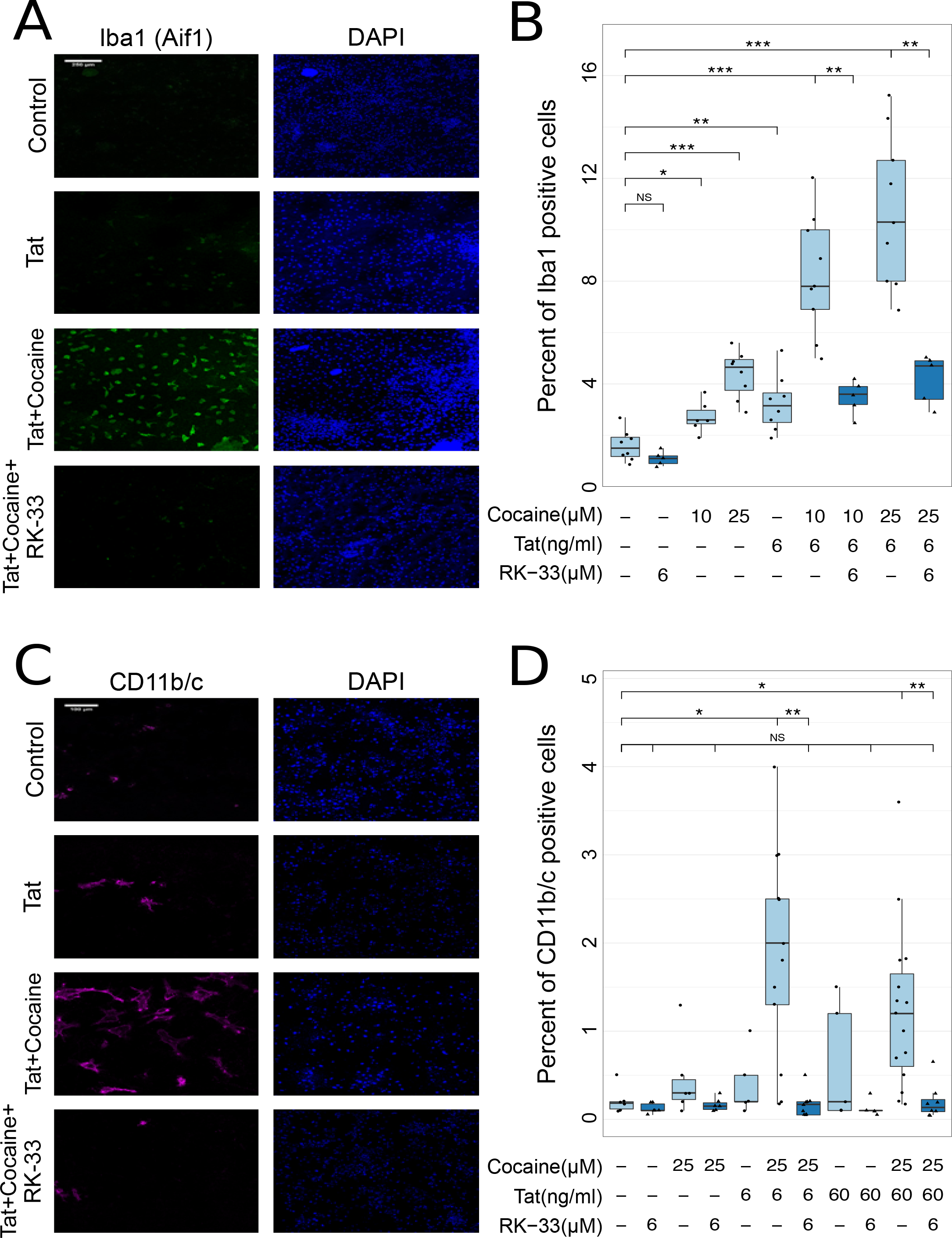
RK-33 treatment suppresses the activation of microglia induced by different combinations of Tat and cocaine on cortical cells. Cortical cultures were treated with Tat alone (6 or 60 ng/ml) or Tat combined with RK-33 (6 μM) for 48 h prior to addition of cocaine (10 or 25 μM) for another 24 h. Cultures were then fixed with 4% PFA, and microglial cells were detected using anti-Iba1 and anti-CD11b/c antibodies as described in Materials and Methods. The number of microglial cells increased when cultured cells were subjected to Tat alone or cocaine alone, and especially, when Tat was combined with cocaine, producing a dramatic increase of activated microglia in the cortical cell cultures (**A**: blue, DAPI; green, Iba1. Scale bar, 250 μm. **C**: blue, DAPI; purple, CD11b/c. Scale bar, 100 μm). The box-and-whisker plot graphs Tat and/or cocaine only treatments in light blue and the addition of RK-33 in dark blue (**B**, **D**). The boxes cover 50% of data in each condition, and the lines within the boxes indicate the median values. The Mann-Whitney-Wilcoxon test was conducted to calculate the statistical significance, followed by Benjamini-Hochberg adjustment of p-values (*, p<0.05; **, p<0.01; ***, p<0.001).

Taken together, the results of the RNA-seq analysis demonstrates that, in agreement with a previous publication (45), Tat triggers the expression of genes associated with activated microglia. Moreover, this activation is inhibited by RK-33. These same genes found by RNA-seq to be regulated by HIV-Tat treatment are also found in the high-ranked genes subset of HAND-associated genes from the MOLIERE analysis (p<0.0025), including Csf1r.

### RK-33 inhibition of the activation of microglia by Tat or combined Tat and cocaine treatment

The hallmarks of microglial activation are the rapid expansion of microglial cells and characteristic changes in cellular morphology (41, 46, 47). Microglial cells were analyzed by immunofluorescent staining of rat cortical cultures using antibodies against two established markers whose expression changes were seen in the RNA-seq data, namely Iba1 and CD11b/c (48, 49). Treatment with RK-33 alone did not affect the number of Iba1 positive cells. Treatment with either Tat (6 ng/ml) or cocaine (10 μM or 25 μM) increased slightly but not significantly the number of Iba1 positive cells, while the combined Tat and cocaine treatment caused a dramatic elevation of Iba1 positive cells (Fig 4A, B, compare light blue (minus RK-33) to dark blue (+RK-33)). The number of Iba1 positive cells in Tat and cocaine co-treated cultures also treated with RK-33 were not significantly different from untreated controls (Fig 3A, B). These results were confirmed with CD11b/c, which is expressed in resting microglia and is greatly elevated upon activation (49, 50). The results were similar to the analysis of Iba1. The combined treatment of Tat and cocaine significantly elevated the number of CD11b/c positive cells, and RK-33 treatment reduced the number of CD11b/c-positive cells that were elevated by Tat/cocaine back to basal untreated control levels (Fig 3C, D, compare light blue (minus RK-33) to dark blue (+RK-33)).

**Figure 4.**
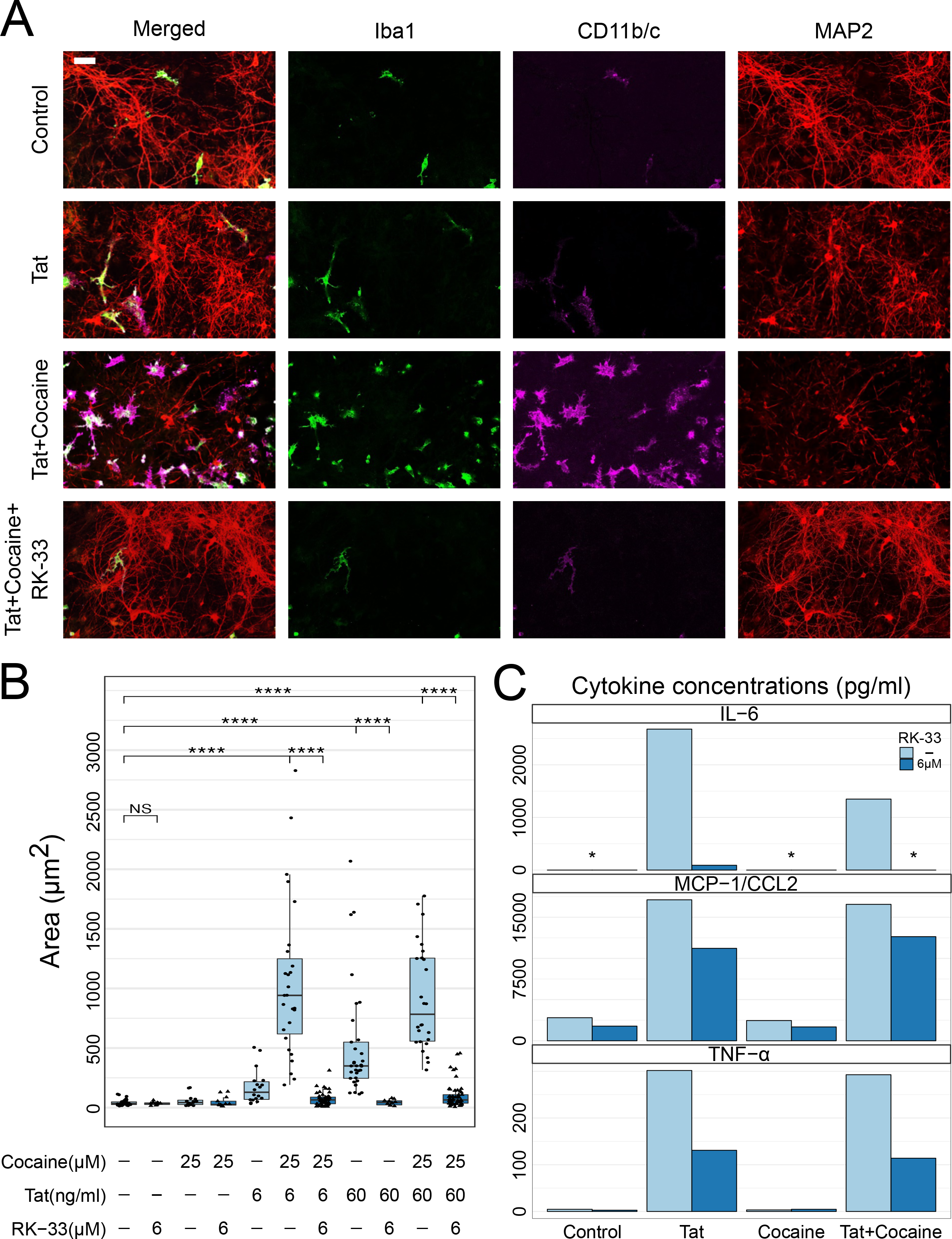
RK-33 treatment decreases the size of microglial cells and attenuates the secretion of proinflammatory cytokines induced by HIV-Tat and cocaine. Cortical cultures were treated with Tat (6 or 60 ng/ml) and/or cocaine (25 μM) alone or Tat combined with RK-33 (6 μM) for 48 h prior to addition of cocaine for another 24 h. Conditioned media were collected from each sample, cultures were then fixed with 4% PFA and microglial cells were detected using anti-Iba1, anti-CD11b/c, and anti-MAP2 antibodies as described in Materials and Methods. The area of microglial cells was quantified using ImageJ (**A**: blue, DAPI; green, Iba1; red, MAP2. Scale bar, 50 μm). The box-and-whisker plot graphs Tat and/or cocaine only treatments in light blue and RK-33 in dark blue (**B**). The boxes cover 50% of data in each condition, and the lines within the boxes indicate the median values. Each point corresponds to an image field, with each image covering a range of 60-100 cells. The Mann-Whitney-Wilcoxon test was conducted to calculate the statistical significance, followed by Benjamini-Hochberg adjustment of p-values (*, p<0.05; **, p<0.01; ***, p<0.001). (**C**) Cortical cultures were treated as above with Tat (60 ng/ml) and/or cocaine (25 μM) alone or Tat combined with RK-33 (6 μM). The cytokine levels were determined by 8-plex chemokine/cytokine array. Bars represent chemokine/cytokine concentrations in the medium (pg/ml). (*) indicates the concentration below the detection limit.

Pathological stimuli are known to trigger morphological remodeling of microglia. Ramified, quiescent microglial cells transition to an intermediate form with less arborization and large soma, followed by amoeboid morphology. The size of microglial cells is greatly increased in the transition (47, 51, 52). Here, the combined Tat and cocaine treatment led to amoeboid morphology of microglial cells detected by immunofluorescence of both CD11b/c and Iba1 (Fig 4A, Supplemental Figure 4A) and increased cell body area (Fig 4A, B). Treatment with RK-33 diminished the morphological and body changes induced by Tat/cocaine treatment (Fig 4A, B, compare light blue (minus RK-33) to dark blue (+RK-33)).

### The secretion of proinflammatory cytokines induced by HIV Tat and cocaine is inhibited by RK-33

The pathological effects of microglia activation are directly associated with the secretion of proinflammatory cytokines. Thus, to measure the level of cytokines we collected media of the cortical cultures treated with Tat, followed by addition of cocaine with or without RK-33. The level of eight cytokines (IL-6, TNF-alpha, MCP1-CCL2, IFN-γ, VEGF, IL-2, IL-4, IL-8, IL-1-α and IL-1-β), was determined with the Luminex Multiplex assay. Tat treatment elevated the concentration of secreted IL-6, TNF-alpha, and MCP1-CCL2, IL-1-α and IL-1-β, while RK-33 treatment abolished the elevation (Fig 4C, Supplemental Figure S4B,C). The concentrations of IFN-γ, VEGF, IL-2, IL-4, and IL-8 were not affected by the treatments (not shown). Surprisingly, Tat and cocaine co-treatment either did not increase or slightly increase the level of cytokines relative to Tat-treated culture, which suggests that the detrimental effects of cocaine involve an additional mechanism.

## Discussion

We recently reported the development of MOLIERE, an AI-based literature mining system to determine target genes and associated small molecules that are potentially useful for testing novel gene-disease connections (23, 24). Here, we applied MOLIERE to uncover novel targets for the treatment of HIV associated neurocognitive disorder. This usage of MOLIERE uncovered DDX3 and its specific inhibitor, RK-33, and we have experimentally verified this novel target as worthy of further investigation. To our knowledge, neither DDX3 nor RK-33 have previously been linked to HAND or other neurodegenerative diseases. RK-33 was developed and tested as an antitumor drug, and importantly in this context, without any indication of systemic toxicity in mice with injections up to 20 mg/kg for seven weeks (29).

The evaluation of RK-33 in a well-established culture model of HAND augmented with a drug of abuse showed dose-dependent inhibition of neuronal apoptosis induced by combined Tat and cocaine treatment with RK-33 concentrations as low as 1 μM. Further, RNA-seq analysis and measurement of secreted pro-inflammatory cytokines demonstrated that microglial activation induced by Tat and cocaine was suppressed by RK-33, thus providing a plausible mechanism for the neuroprotective effects downstream of DDX3 inhibition. The dual role of microglial cells in HAND pathogenesis as a brain reservoir for viral replication and source of secreted viral proteins, and as a modulator of the inflammatory response is well established (53, 54). Our analysis confirmed previous results showing an activation of microglia by Tat and a synergistic effect of cocaine (35, 54, 55), as measured by elevation in the number of Iba1 and CD11b/c positive cells in cortical cultures and the induction of morphological changes and enlargement of microglial cells.

Collectively, these results are the first to connect DDX3 to the activation of microglia. Moreover, the DDX3 inhibitor RK-33 inhibits Tat/cocaine-dependent microglial activation, which both implicates DDX3 in this pathogenesis and also suggests it is a viable target for treatment of HAND and potentially other neurocognitive diseases in which activated microglia play a role. The mechanism of DDX3-dependent regulation of microglia activation remains to be determined. However, recent results regarding the function of DDX3 in the regulation of a macrophage inflammatory response may shed light on DDX3 activity in neuroinflammation. HIV proteins activate the secretion of chemokines and cytokines by microglia cells through NF-κB, p38 and TGF-β pathways (56–59). The small GTPase Rac1 was shown to be a regulator activating morphological changes in microglia cells (60, 61). These pathways are well known to control the inflammatory response, cytokine secretion, and migration of macrophages. Importantly, it was recently shown that DDX3 directly regulates the translation of p38 MAPK, Rac1, STAT1 (TGF-β) and TAK1, which play essential roles in NF-κB regulation (38). Direct roles of DDX3 in regulating pro-inflammatory responses in the pathogenesis of bowel disease and Listeria infection were also recently observed (62, 63). Inhibition of DDX3 activity stalled the translation of target proteins (38, 64), resulting in a decrease of cytokine secretion and inhibition of macrophage migration and phagocytosis (38). It therefore appears plausible that DDX3-dependent translational control may be the mechanism that regulates microglial activation in neuroinflammatory pathways. Experiments are ongoing to formally test this.

As mentioned previously, DDX3 has been proposed as a target for development of anti-viral and anti-HIV therapy (65, 66) (67–70). This makes DDX3 a unique target, in which inhibition may affect both HIV-related neurotoxicity and the production of viral proteins by glial cells. However, the contribution of DDX3 to innate immunity has to be fully evaluated prior to clinical advancement of DDX3 inhibitors for HAND therapy (27, 71). Nonetheless, DDX3-specific inhibition as a target and RK-33 as a prototype molecule for the development of HAND therapy has been validated and should be investigated further. DDX3X mutations have been recently connected to female intellectual disability (72), a developmental syndrome linked to the effect of DDX3X on migration and differentiation of neuronal progenitors (73) and neurite development (74). Although these apparently seem linked to the role of DDX3 during early development, the latter may cause potential negative effects of DDX3-specific therapy on brain development and needs to be carefully evaluated.

To conclude, given the importance of microglial activation in the pathology of other neurodegenerative diseases, DDX3 targeting could be applicable for the treatment of many neurodegenerative diseases.

## Materials and Methods

### MOLIERE Analysis

The implementation and documentation are available at https://github.com/JSybrandt/MOLIERE. The repository also contains list of all software dependencies to packages to compute approximate nearest neighbor graphs, low-dimensional embeddings, probabilistic topic modeling, phrase mining, and graph algorithms. The repository is organized in two major sub-projects, namely, *build_network* and *run_query*, each contains its own documentation. Preinstallation dependencies include gcc 5.4 or greater, Python 3, Java 1.8, Scons, Google Test, and Mpich 3.1.4. It is recommended to use parallel machines, as many components of the project are parallelized being too slow if executed in sequential mode. The input to the phase of building the knowledge network requires downloading full MEDLINE and UMLS. Building the network is also possible with partial MEDLINE if one wants to restrict the information domain in order to increase speed. Most algorithmic components require parameters that are provided with the code. When the knowledge network is constructed, the second phase consists of running queries using run query subproject. Running all queries, each of type gene-HAND, can be done in parallel, as all of them are independent. Each query will return the hypotheses in a form of topic model, i.e., a distribution of most representative keywords per learned topic, as well as a ranking score. In addition, the result of each single query can be visualized for further analysis using the *visualization* sub-project that can be found in the same repository. The visualization connects all learned topics in a network, where nodes correspond to topics, and edges represent mutual content connections. Clicking each node will bring up the most relevant to the corresponding paper topics as well as the most representative topical keywords.

Here we describe the datasets used in knowledge network construction and querying MEDLINE, the NLM database of biomedical abstracts, releases public yearly baselines. We used the 2017 baseline, published early that year, which was the most up-to-date at the time. This dataset consists of 26,759,399 documents; however, we found that certain short documents hinder hypothesis generation results. Therefore, we removed any document that is fewer than 20 words that also does not contain at least two “rare” words found in the bottom 85% most frequent words. The Unified Medical Language System (UMLS) consists of known medical entities, as well as their synonyms and relationships. This NLM dataset releases every six months, and we used the “2017AB” release, also the most recent available to us at the time of our experiments. This release consists of 3,639,525 entities. SemMedDB, another dataset produced by the NLM, which contains automatically extracted predicates from MEDLINE documents and keeps a six-month release schedule. We downloaded the December 31^st^, 2017 release consisting of 15,836,301 unique subject-verb-object statements, as well as corresponding UMLS types and MEDLINE identifiers. Lastly, the HUGO gene dataset collects human gene symbols. Unlike the NLM sources, HUGO follows rolling updates and does not keep numbered versions. We leveraged the “complete HGNC dataset” from January 19^th^, 2018, which contained 42,139 gene symbols. From this initial set, we filtered out 1,248 symbols that could not be found in either our MEDLINE subset or our UMLS release, as our system has no known information on these gene symbols. BindingDB (31) publishes protein-ligand structures associated with gene symbols. This dataset additionally supplies rolling releases, which we accessed on January 8^th^, 2018. This dataset contains 1,507,528 binding measurements, within which we identified 1,202 human gene symbols from HUGO (30) under the field “Entry Name of Target Chain.” For each gene, we recorded both the total number of dataset occurrences as well as the distinct names found under the field “Target Name Assigned by Curator or DataSource.”

### RNA sequencing and analysis

RNA and library preparation, post-processing of the raw data and data analysis were performed by the USC CTT COBRE Functional genomics Core. RNAs were extracted with Qiagen RNeasy Mini kit as per manufacturer recommendations (Qiagen, Valencia, CA, USA) and RNA quality was evaluated on RNA-1000 chip using Bioanalyzer (Agilent, Santa Clara, CA, USA). RNA libraries were prepared using an established protocol with NEBNExt Ultra II Directional Library Prep Kit (NEB, Lynn, MA). Each library was made with one of the TruSeq barcode index sequences, and the Illumina sequencing was performed by GENEWIZ, Inc. (South Plainfield, NJ) with Illumina HiSeq4000 (150bp, pair-ended). Sequences were aligned to the Mus Musculus genome GRCm38.p5 (GCA_000001635.7, ensemble release-88) using STAR v2.4 (75). Samtools (v1.2) was used to convert aligned sam files to bam files, and reads were counted using the featureCounts function of the Subreads package (76) with Gencode.vM19.basic.annotation.gtf annotation file. Only reads that were mapped uniquely to the genome were used for gene expression analysis. Differential expression analysis was performed in R using the edgeR package (77). The average read depth for the samples was around 15 million reads, and only genes with at least one count per million average depth were considered for differential expression analysis. Raw counts were normalized using the Trimmed Mean of M-values (TMM) method. The normalized read counts were fitted to a quasi-likelihood negative binomial generalized log-linear model using the function glmQLFit, and genewise statistical tests for significant differential expression were conducted with empirical Bayes quasi-likelihood F-tests using the function glmQLFTest.

#### Pathway enrichment and GSEA analysis

Pathway enrichment of HIV-Tat and RK-33 regulated genes were analyzed using Enrichr (78) R package (https://CRAN.R-project.org/package=enrichR) with GO, KEGGs pathway and EnrichrPathway databases. The GSEA enrichment analysis were performed with Broad Institute software (43) (http://software.broadinstitute.org/gsea/index.jsp) with custom MSigDB, represented genes enriched in microglia, neurons and astrocytes. The MSiDB were built as described in GSEA documentation (https://software.broadinstitute.org/gsea/doc/GSEAUserGuideFrame.html). The gene lists were prepared from Zhang dataset of transcriptome profiling of glia neurons and vascular cells of the cerebral cortex (44), https://web.stanford.edu/group/barres_lab/brain_rnaseq.html. The genes which expressed 10 or more fold higher in one cell type relative to any other had been assigned as cell-type specific (Supplemental Table S3). Overrepresentation of Tat-activated genes in MOLIERE-selected subset were identified using hypergeometric distribution test.

#### Preparation and cultivation of primary cortical cultures

Primary cortical cell cultures were prepared from 18-day-old Sprague-Dawley (Envigo Laboratories, Indianapolis, IN) rat fetuses or from 18-day-old C57BL/6 mouse fetuses as previously described (79, 80). Procedures were carried out in accordance with the University of South Carolina Institutional Animal Care and Use Committee. Briefly, cortical regions were dissected and incubated for 10 min in a solution of 0.05% Trypsin/EDTA in Hank's balanced salt solution (HBSS) (Thermo Fisher Scientific). The tissue was then exposed for 5 min to soybean trypsin inhibitor (1 mg/ml in HBSS) and rinsed 3 times in HBSS. Cells were dissociated by trituration and distributed to poly-L-lysine coated 12-well plates with inserted round glass coverslips. Alternatively, cortical cell cultures were grown in 6-well plastic plates (VWR International, Radnor, PA).

At the time of plating, plates contained DMEM/F12 (Thermo Fisher Scientific) supplemented with 100 mL/L fetal bovine serum (HyClone, Thermo Fisher Scientific). After a 24-hr period, DMEM/F12 was replaced with an equal amount of serum-free Neurobasal medium supplemented with 2% v/v B-27, 2 mM GlutaMAX supplement and 0.5% w/v D-glucose (all reagents from Thermo Fisher Scientific). Half of the Neurobasal medium was replaced with freshly prepared medium of the same composition once a week. Cultures were used for experiments at the age of 10-12 days in vitro (DIV).

### Experimental treatments

Primary cortical cultures were treated with recombinant Tat 1-86 (Diatheva, Italy), at concentrations ranging from 5 ng/ml up to 60 ng/ml as described in Figures and Figure legends. The concentrations of Tat had been selected to represent the estimated Tat concentration in the brain of HIV-infected patients (81). Cocaine-HCl was obtained from Sigma Chemicals and was dissolved in sterile water immediately before the addition to cell cultures. Cocaine concentrations ranged from 10 μM up to 1000 μM as described in Figures. The concentration of cocaine is in a range of the brain cocaine concentration estimated for recreational users based on animal studies (82) and postmortem examination of the brain tissues in fatal cases of cocaine abuse (83). RK-33, a small molecule inhibitor of DDX3, was obtained from Selleck Chemicals, (Catalog No.S8246, Selleck Chemicals, Houston, Texas). A stock solution of RK-33 was prepared in DMSO (5 mM) and was diluted to final concentrations from 0.25 μM up to 12 μM.

### Apoptotic/Dead cells detection

Dead and apoptotic cells were detected using CellEvent Caspase-3/7 Kit (#C10423, Thermo Fisher Scientific) according to the manufacturer’s recommendations. Briefly, after experimental treatment, Caspase3/7 Green Detection Reagent was added directly to cells, and the plate was incubated 30 min at 37°C. The final concentration of the reagent was 500 nM. During the final 5 min of incubation, SYTOX AADvanced dead cell solution was added. The final concentration of the stain was 1 μM. Cells were rinsed with PBS, and images of live cells were taken immediately. Alternatively, cells were fixed with 4% paraformaldehyde, imaged, and used for further experiments.

### Immunocytochemistry

For ICC analysis cells were plated on glass coverslips and placed inside 12-well plates. Following experimental treatment, primary neuronal cultures were fixed with 4% paraformaldehyde and permeabilized with 0.1% Triton X-100. Fixed cultures were blocked with 10% fetal bovine serum for 2 hours and then co-labeled overnight with different primary antibodies: chicken polyclonal anti-MAP2 antibodies (1:2,500) (# ab92434, Abcam, Cambridge MA), rabbit monoclonal anti-Iba1 (1:200) (# ab178847 Abcam, Cambridge MA), human recombinant anti-CD11b/c (1:50) (#130-120-288, Miltenyi Biotec, Germany). Secondary antibodies, goat anti-chicken IgG conjugated with AlexaFluor 594, and goat anti-mouse IgG conjugated with AlexaFluor 488 (1:500; Invitrogen Life Technologies, Grand Island NY), were used for visualization. Anti-CD11b/c antibodies were originally labeled with allophycocyanine. To identify cell nuclei, DAPI was added with the final PBS wash, and coverslips were mounted on glass slides using VECTASHIELD Vibrance mounting medium (Vector Laboratories, Burlingame, CA).

### Image processing and analysis

Images were taken on a Carl Zeiss LSM 700 laser scanning confocal microscope (Carl Zeiss Meditec, Dublin, CA) equipped with 20x (Plan APO 0.8 air) or 63x (Plan APO 1.4 oil DIC) objectives. Images were captured using 1.0 scanning zoom with 312-nm (20x) or 142-nm (63x) X-Y pixel size. Fluorescence and differential interference contrast (DIC) imaging was done using single-frame or tile (3×3 or 3×4) modes.

ImageJ software (National Institutes of Health, USA) was used for manual or automatic analysis of microscopy images acquired using a Zeiss 700 confocal microscope. Total number of cells and percentage of Iba1 or CD11b/c - positive cells were estimated using segmentation of DNA channel (DAPI) followed by “Analyze Particles” ImageJ command. Size of microglia cells was estimated individually using “Freehand selections” ImageJ tool. Data were aggregated, analyzed and visualized using R gglot2 tools.

Background correction of widefield images was performed by background (Gaussian blur) division procedure (32-bit mode) followed by image histogram adjustment for 16-bit dynamic range.

### Cytokine/Chemokine Array

Cortical cultures medium was collected, frozen and sent to Eve Technologies Corporation (Calgary, Canada) for the LUMINEX based analysis of cytokines by Featured-Rat Cytokine Array/ Chemokine Array 8-plex (RECYT-08-204) and 27-plex (RD27). The 8-plex array analyzed IFN-γ, IL-2, IL-4, IL-6, IL-18, MCP-1, TNF-α, and VEGF. The 27-plex array analyzed Eotaxin, EGF, Fractalkine, IFN-gamma,IL-1alpha, IL-1beta, IL-2, IL-4, IL-5, IL-6, IL-10, IL-12(p70), IL-13, IL-17A, IL-18, IP-10, GRO/KC, TNF-alpha, G-CSF, GM-CSF, MCP-1, Leptin, LIX, MIP-1alpha, MIP-2, RANTES, VEGF

## Acknowledgments

We thank Dr Amar N. Kar for the help with primary cortical cultures, Drs. Jeffery L. Twiss, Anna Kashina, Pavel Ortinski and Inna Grosheva for fruitful discussions and critical reading of the manuscript. We than the cores of COBRE Center for Targeted Therapeutics for transcriptomics analysis (Functional Genomics Core) and microscopy and image analysis (Microscopy and Flow cytometry Core). We thank Drs Chinenov and Oliver (The David Z. Rosensweig Genomics Research Center, HSS, NY) for consultation and help with data visualization. The work was supported by awards from NIH NIDA R21DA047936 and R03DA043428 (MS), R01DA035714 (JZ), NIH CA223956 (MW).

## Contributions

MS, IS, MDW, JZ initiated the study, JS and IS developed software and performed literature mining, MA, CB, DO, ML, MS, MDW designed and performed experiments, CB and EP performed statistical analysis, HJ and MS performed bioinformatics analysis, VS helped with image acquisition and image analysis, JRT, SL, EB and JZ helped with data analysis and provided critical suggestions, MA, MDW, IS and MS wrote the manuscript.

## Competing interests

The authors declare no competing interests.

