## Supplemental Figures and Figure legends for "Inhibition of the Dead Box RNA Helicase 3 prevents HIV-1 Tat and cocaine-induced neurotoxicity by targeting microglia activation"

#### Supplemental Figure S1. RK-33 protects neurons in primary mouse cortical cultures from the combined neurotoxicity of Tat and cocaine

Primary mouse cortical cultures were treated with Tat (60 ng/ml) and/or cocaine (500 µM) alone or Tat combined with a range of RK-33 for 48 h prior to addition of cocaine for another 24h. Cultures were then fixed with 4% PFA and the level of cell perturbation was detected using CellEvent Caspase 3/7 Assay Kit (**A**: green, Caspase 3/7; red, Ethidium Bromide).

Tat and/or cocaine only treatments are colored with green, and the deepening blue hue represents the increasing concentration of RK-33 (2,4, and 6 µM) (**B**).

The bar heights mark the mean values and the error bars cover one sample standard error from the sample mean. Each point corresponds to an image, with each image containing a range of 60-100 cells. The Mann-Whitney-Wilcoxon test is conducted to calculate the statistical significance, followed by Benjamini-Hochberg adjustment of p-values (*, $p<0.05$; **, $p<0.01$; ***, $p<0.001$).

#### Supplemental Figure S2. Pathway enrichment analysis of Tat and RK-33 regulated genes.

Functional enrichment analysis with Enrichr of genes regulated by Tat relative to control, and Tat plus RK-33 relative to Tat (adjusted FDR $p<0.05$). Color represents the adjusted p value of the enrichment and the gene ratio of enriched pathway/GO-category in the datasets. **(A)** Enrichment of GO (Gene Ontology)- Molecular function category, **(B)** Enrichment of GO-Biological processes category, **(C)** Enrichment of genes associated with specific pathways.

#### Supplemental Figure S3. Gene set enrichment analysis of brain cell-type specific genes of Tat and RK-33 regulated genes.

(**A**) GSEA plots of the distribution of microglia, neurons and astrocytes – specific genes in Tat relative to control and Tat plus RK-33 relative to Tat datasets. The microglia-specific genes mostly associated with Tat upregulated genes (upper panel) and RK-33 downregulated genes (lower panel). (**B**) Metrics of GSEA analysis results. Absolute value of normalized enrichment score (NES) is the highest for microglia associated genes in both datasets.

#### Supplemental Figure S4. RK-33 treatment reverses the changes in the morphology of microglial cells induced by Tat and cocaine and inhibits cytokine production

**(A)** Cortical cultures were treated with Tat (6 ng/ml) and/or cocaine (25 µM) alone or Tat combined with 6 µM of RK-33 for 48 h prior to addition of cocaine for another 24 h. Cultures were then fixed with 4% PFA and the changes in microglial cell morphology were detected using anti-Iba1 and anti-CD11b/c antibodies. (green, Iba1; purple, CD11 b/c). Scale bar, 25 µm.

**(B)** Cortical cultures were treated as above with Tat (60 ng/ml) and/or cocaine (25 µM) alone or Tat combined with RK-33 (6 µM). The cytokine levels were determined by 27-plex chemokine/cytokine array. Bars represent chemokine/cytokine concentrations in the medium (pg/ml). (*) indicates the concentration below the detection limit.

(**C**) Cortical cultures were threated with Tat (60 ng /ml) alone or Tat combined with 6 µM of RK-33 for 72 h prior to addition of cocaine for another 24 h. Samples of culture medium were collected at the beginning of the treatment (0 hour), at 72 h of treatment (before the addition of cocaine) and at 96 h of treatment (24 h of combined treatment). The cytokines level had been determined by 8-plex chemokine/cytokine array. Bars represent chemokine/cytokine concentrations in the medium (pg/ml). (*) indicates the concentration below detection limit.

### Supplemental Tables

Supplemental Table S1. MOLIERE output of the genes associated with HAND

Supplemental Table S2. RNA-seq analysis of transcriptome regulation by Tat and RK-33 in the cortical cultures

Supplemental Table S3. Brain cell-type specific genes used for GSEA.

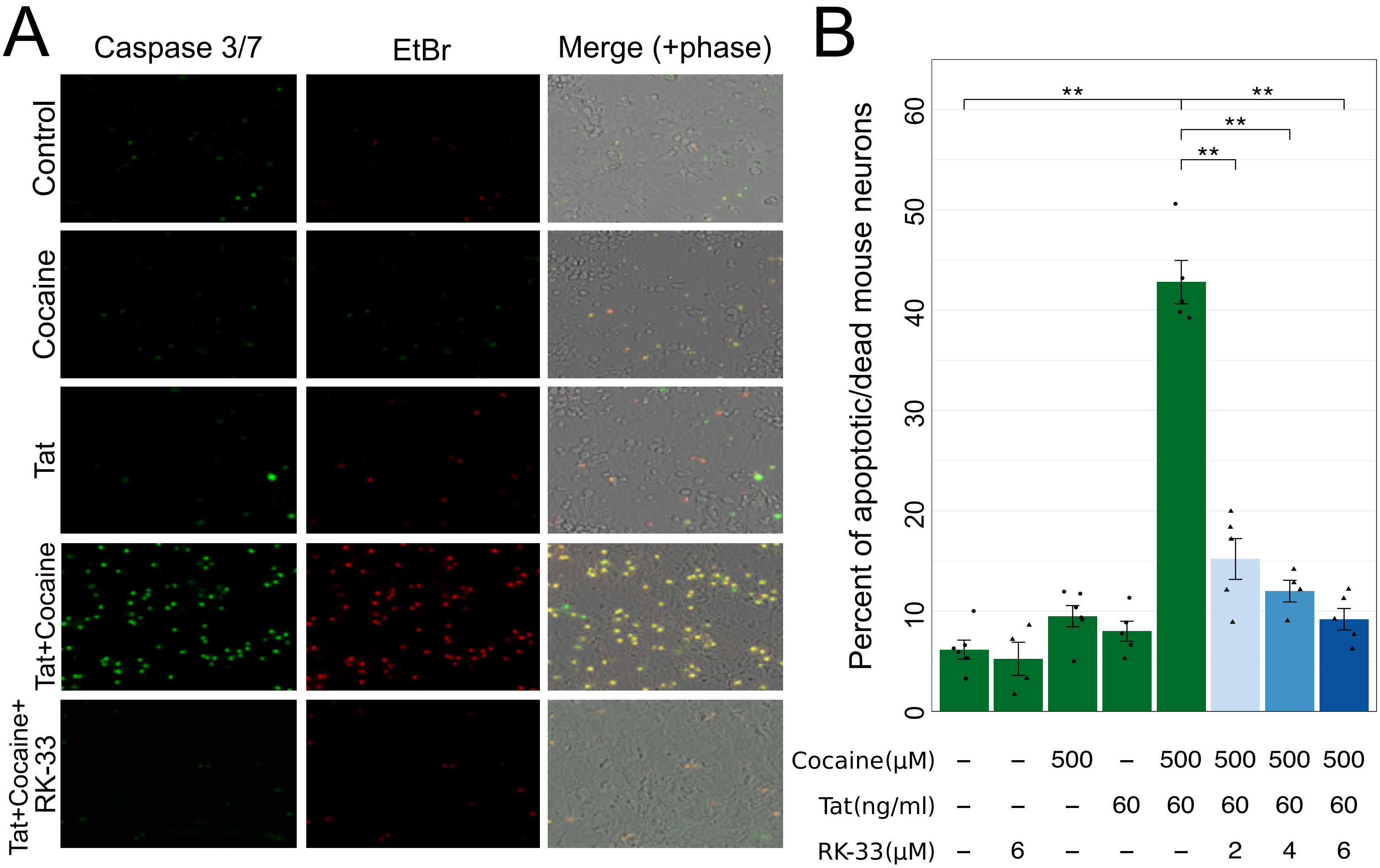

A
Caspase 3/7
EtBr
Merge (+phase)
Control
Cocaine
Tat
Tat+Cocaine
Tat+Cocaine+
RK-33
B
Percent of apoptotic/dead mouse neurons
0
10
20
30
40
50
60
**
**
**
**
Cocaine(μM) – – 500 – 500 500 500 500
Tat(ng/ml) – – – 60 60 60 60 60
RK-33(μM) – 6 – – – 2 4 6

A

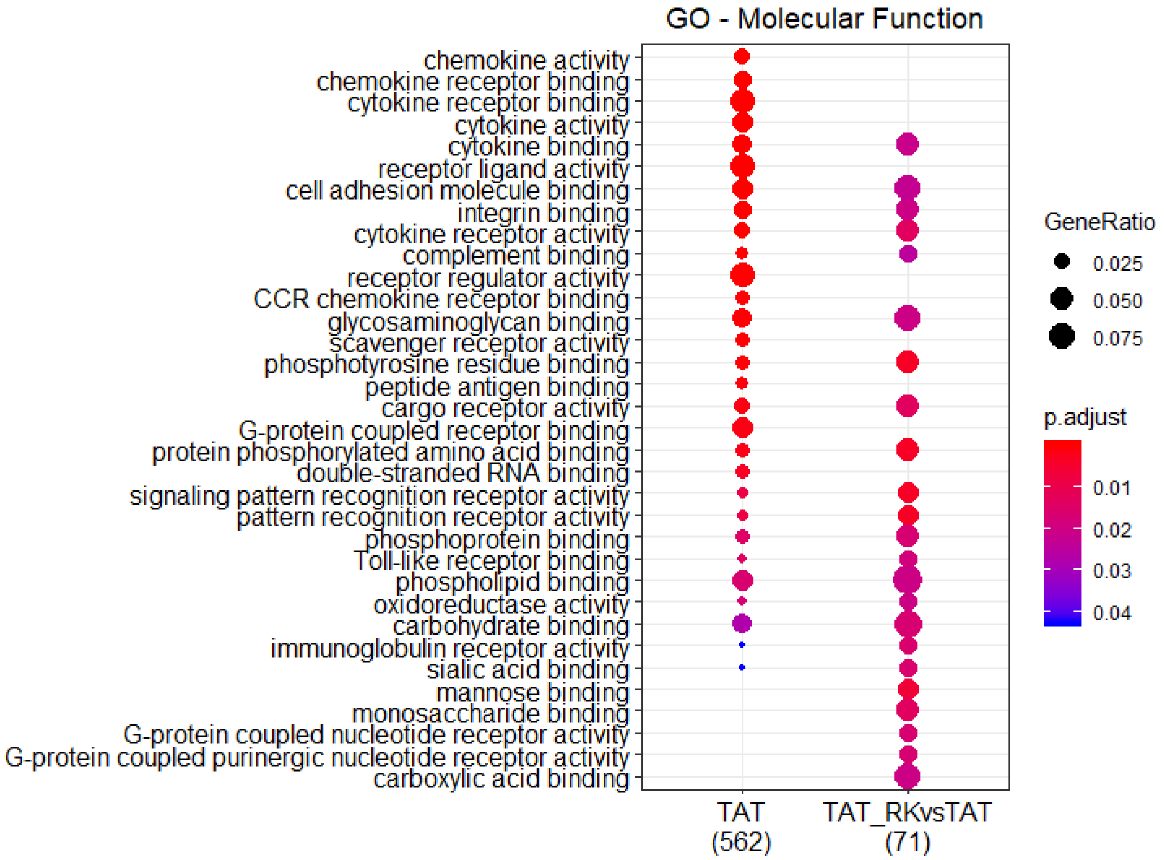
GO - Molecular Function
chemokine activity
chemokine receptor binding
cytokine receptor binding
cytokine activity
cytokine binding
receptor ligand activity
cell adhesion molecule binding
integrin binding
cytokine receptor activity
complement binding
receptor regulator activity
CCR chemokine receptor binding
glycosaminoglycan binding
scavenger receptor activity
phosphotyrosine residue binding
peptide antigen binding
cargo receptor activity
G-protein coupled receptor binding
protein phosphorylated amino acid binding
double-stranded RNA binding
signaling pattern recognition receptor activity
pattern recognition receptor activity
phosphoprotein binding
Toll-like receptor binding
phospholipid binding
oxidoreductase activity
carbohydrate binding
immunoglobulin receptor activity
sialic acid binding
mannose binding
monosaccharide binding
G-protein coupled nucleotide receptor activity
G-protein coupled purinergic nucleotide receptor activity
carboxylic acid binding
TAT (562)
TAT_RKvsTAT (71)
GeneRatio
0.025
0.050
0.075
p.adjust
0.01
0.02
0.03
0.04

B

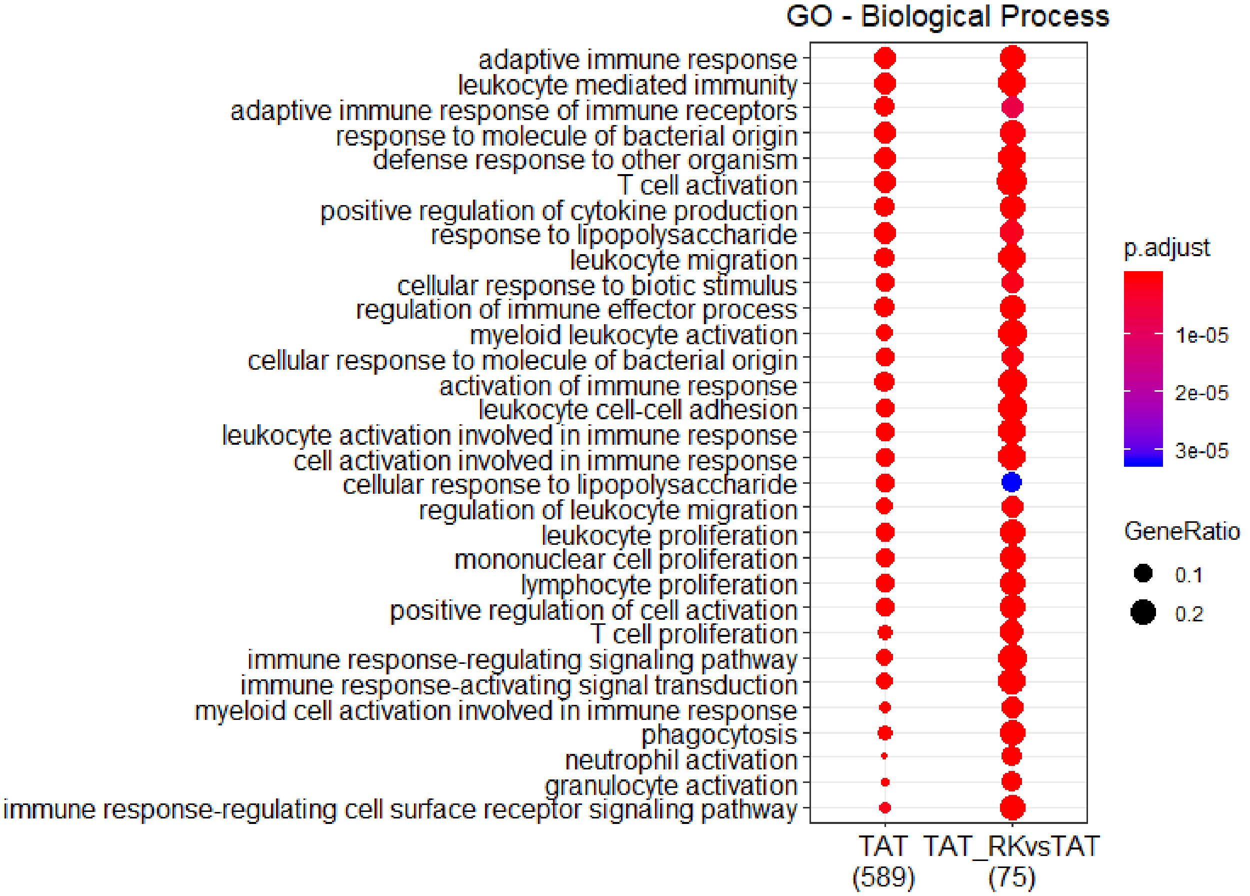
GO - Biological Process
adaptive immune response
leukocyte mediated immunity
adaptive immune response of immune receptors
response to molecule of bacterial origin
defense response to other organism
T cell activation
positive regulation of cytokine production
response to lipopolysaccharide
leukocyte migration
cellular response to biotic stimulus
regulation of immune effector process
myeloid leukocyte activation
cellular response to molecule of bacterial origin
activation of immune response
leukocyte cell-cell adhesion
leukocyte activation involved in immune response
cell activation involved in immune response
cellular response to lipopolysaccharide
regulation of leukocyte migration
leukocyte proliferation
mononuclear cell proliferation
lymphocyte proliferation
positive regulation of cell activation
T cell proliferation
immune response-regulating signaling pathway
immune response-activating signal transduction
myeloid cell activation involved in immune response
phagocytosis
neutrophil activation
granulocyte activation
immune response-regulating cell surface receptor signaling pathway
TAT (589)
TAT_RKvsTAT (75)
p.adjust
1e-05
2e-05
3e-05
GeneRatio
0.1
0.2

C

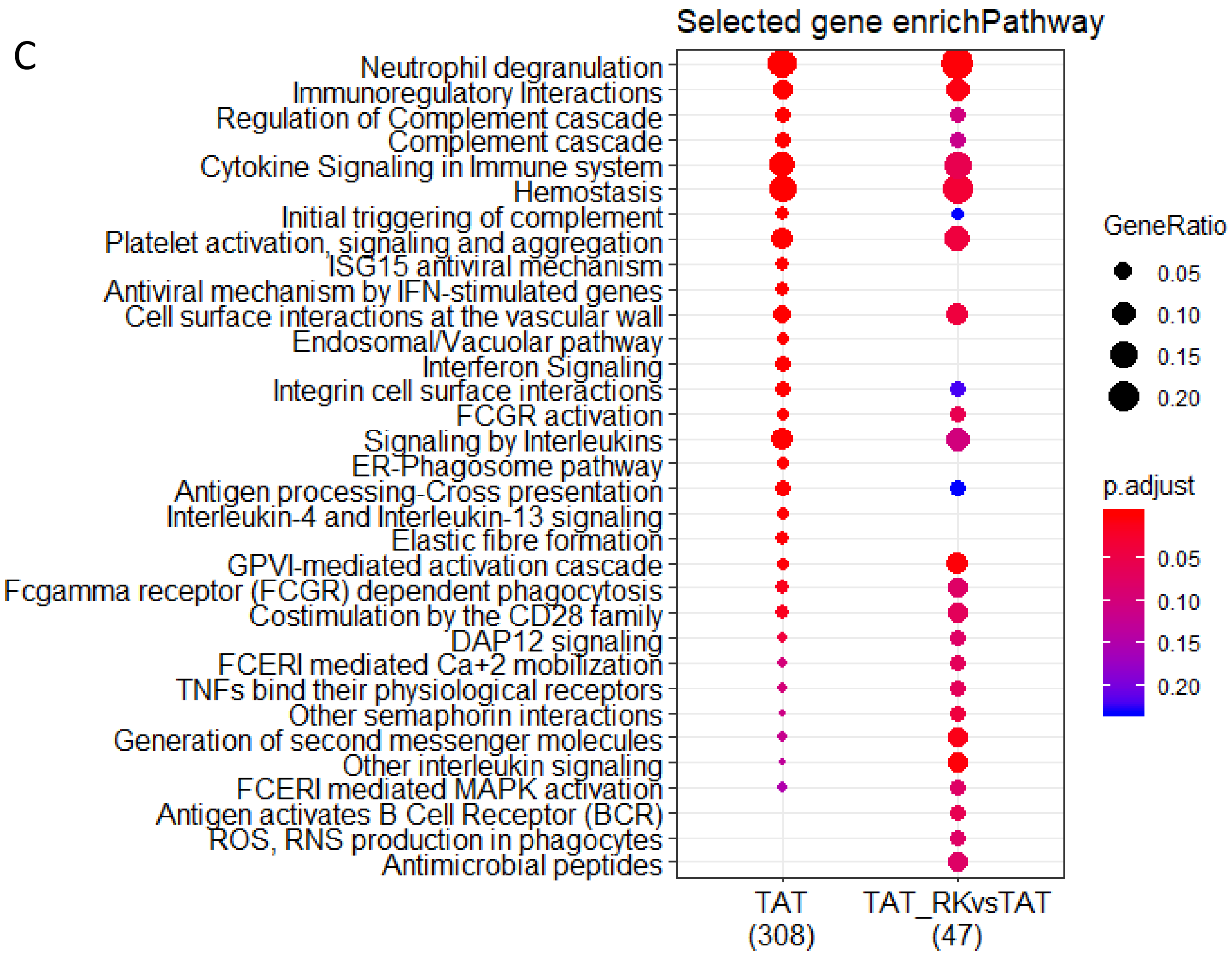
Selected gene enrichPathway
Neutrophil degranulation
Immunoregulatory Interactions
Regulation of Complement cascade
Complement cascade
Cytokine Signaling in Immune system
Hemostasis
Initial triggering of complement
Platelet activation, signaling and aggregation
ISG15 antiviral mechanism
Antiviral mechanism by IFN-stimulated genes
Cell surface interactions at the vascular wall
Endosomal/Vacuolar pathway
Interferon Signaling
Integrin cell surface interactions
FCGR activation
Signaling by Interleukins
ER-Phagosome pathway
Antigen processing-Cross presentation
Interleukin-4 and Interleukin-13 signaling
Elastic fibre formation
GPVI-mediated activation cascade
Fcgamma receptor (FCGR) dependent phagocytosis
Costimulation by the CD28 family
DAP12 signaling
FCERI mediated Ca+2 mobilization
TNFs bind their physiological receptors
Other semaphorin interactions
Generation of second messenger molecules
Other interleukin signaling
FCERI mediated MAPK activation
Antigen activates B Cell Receptor (BCR)
ROS, RNS production in phagocytes
Antimicrobial peptides
TAT (308)
TAT_RKvsTAT (47)
GeneRatio
0.05
0.10
0.15
0.20
p.adjust
0.05
0.10
0.15
0.20

A

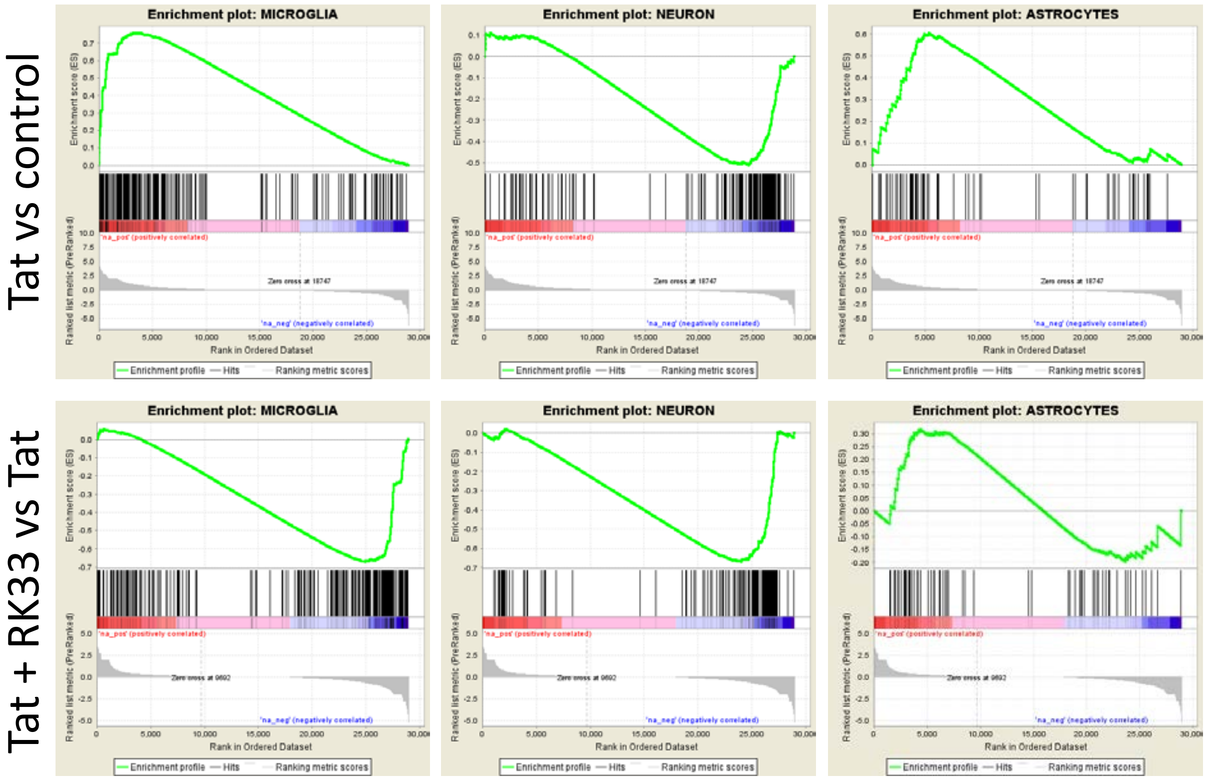


B

| | HIV-Tat vs control | | | Tat + RK33 vs Tat | | |
|---|---|---|---|---|---|---|
| GeneSet | MICROGLIA | NEURON | ASTROCYTES | MICROGLIA | NEURON | ASTROCYTES |
| Enrichment Score (ES) | 0.7579531 | -0.512294 | 0.6048508 | -0.67033744 | -0.6691555 | 0.31654462 |
| Normalized Enrichment Score (NES) | 2.2569964 | -1.676707 | 1.543905 | -1.8730992 | -1.7906189 | 0.8456011 |
| Nominal p-value | 0 | 0 | 0.001436782 | 0 | 0 | 0.8576271 |
| FDR q-value | 0 | 0 | 2.51E-04 | 0 | 0 | 0.9289215 |
| FWER p-Value | 0 | 0 | 0.001 | 0 | 0 | 0.511 |

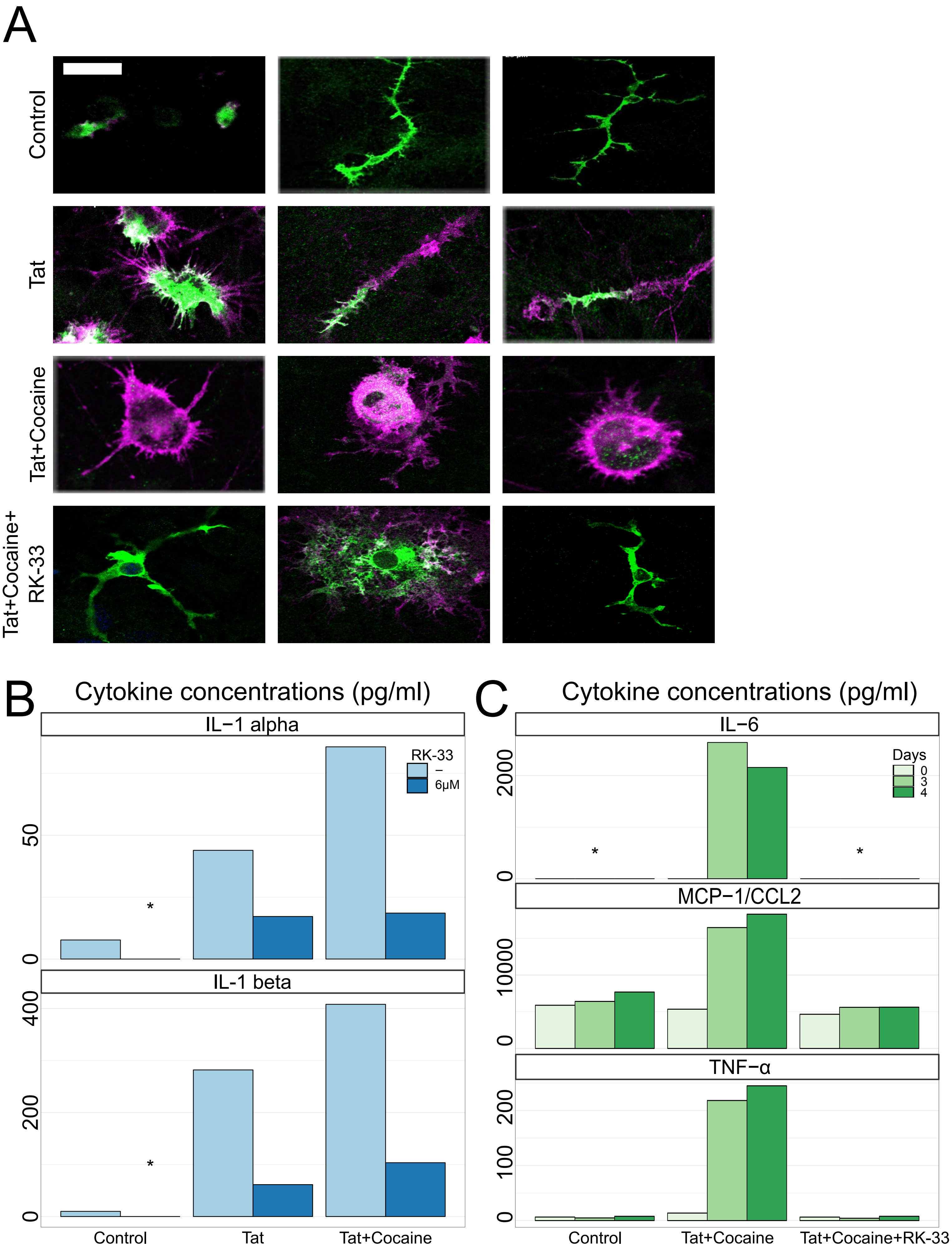

A
Control
Tat
Tat+Cocaine
Tat+Cocaine+ RK-33
B
Cytokine concentrations (pg/ml)
IL−1 alpha
RK-33
−
6µM
50
0
*
IL−1 beta
400
200
0
*
Control
Tat
Tat+Cocaine
C
Cytokine concentrations (pg/ml)
IL−6
Days
0
3
4
2000
0
*
*
MCP−1/CCL2
10000
0
TNF−α
200
100
0
Control
Tat+Cocaine
Tat+Cocaine+RK-33
